# Interpersonal Neural Synchrony Across Levels of Interpersonal Closeness and Social Interactivity

**DOI:** 10.1101/2025.03.18.643871

**Authors:** Alessandro Carollo, Andrea Bizzego, Verena Schäfer, Carolina Pletti, Stefanie Hoehl, Gianluca Esposito

**Author notes:** **Corresponding Author** *Email address:* (Gianluca Esposito).

## Abstract

Interpersonal neural synchrony is a fundamental aspect of social interactions, offering insights into the neural mechanisms underlying human connection and developmental outcomes. So far, hyperscanning studies have examined synchrony across different dyads and tasks, leading to inconsistencies in experimental findings and limiting cross-study comparability. This variability has posed challenges for building a unified theoretical framework for neural synchrony. This study investigated the effects of interpersonal closeness and social interactivity on neural synchrony using functional near-infrared spectroscopy hyperscanning. We recorded brain activity from 142 dyads (70 close-friend, 39 romantic-partner, and 33 mother-child dyads) across three interaction conditions: video co-exposure (passive), a cooperative game (structured active), and free interaction (unstructured active). Neural synchrony was computed between participants’ bilateral inferior frontal gyrus (IFG) and temporoparietal junction (TPJ) using wavelet transform coherence. Results showed that true dyads exhibited significantly higher synchrony than noninteracting surrogate dyads (*q*s <.001, Cohen’s *d* range: 0.17-0.32), particularly in combinations involving the right IFG. Mother-child dyads displayed lower synchrony than adult-adult dyads at the network (*p* <.001) and local level of analysis, pointing to possible developmental and maturational influences on neural synchrony. At the network level, synchrony was highest during video co-exposure, followed by the cooperative game and free interaction (*p* <.001). However, left IFG–left IFG and left IFG–right TPJ synchrony peaked during the cooperative game. Although these effects were statistically significant, the overall impact of social interactivity on interpersonal neural synchrony was small, suggesting that the complexity and richness of social exchanges alone may only modestly influence neural synchrony in naturalistic contexts. By comparing different types of dyads and interaction contexts, this study highlights factors that may guide future hypothesis-driven hyperscanning research and contribute incremental evidence to ongoing efforts to understand the neural mechanisms underlying human social interactions.

## 1. Introduction

“I feel like somehow our hearts have become intertwined. Like when she feels something, my heart moves in tandem. Like we are two boats tied together with a rope. Even if you want to cut the rope, there is no knife sharp enough to do it.” (H. Murakami, *Men Without Women*)

Social interactions play a central role throughout human life. From birth, early social interactions scaffold the acquisition of cognitive, emotional, and social skills (Vygotsky, 1978), including language (Kuhl, 2007, 2011), emotional regulation (Bornstein and Esposito, 2023; Hollenstein et al., 2017), and expectations about the social world that allow engaging in effective interactions with significant others throughout life (Bowlby and Holmes, 2012).

Due to the central role of social interactions in human life, understanding and scientifically characterizing their quality has become a key focus in developmental and social research. A growing body of literature in human development highlights three foundational dimensions of social exchanges: rhythm, reciprocity, and synchrony (e.g., Feldman, 2007; Stern, 2009; Trevarthen, 1998). Rhythm refers to the adjustment of the temporal structure and pacing of interactive behaviors, such as the alternation of vocalizations or gestures between partners, in order to create temporal regularities that facilitate the interaction (De Reus et al., 2021). For instance, caregivers often offer rhythmic stimulation to their infants through activities such as singing, patting, or rocking, which supports the infant’s ability to regulate physiological and emotional states (De Reus et al., 2021; Trehub et al., 2015). Similarly, in adult interactions, social partners adapt their speech rhythms, such as speech rate and prosody, to one another, facilitating accurate timing predictions and stimulus processing (Abney et al., 2014; De Reus et al., 2021; Haegens and Golumbic, 2018; Wilson and Wilson, 2005). Reciprocity refers to the mutual and contingent responsiveness between interacting individuals, where each partner’s behavior not only influences but is also influenced by the other’s response (Anderson et al., 1977; Apicella et al., 2013). It is reflected in the capacity and motivation to engage in joint action and to share psychological states, such as intention, or emotion, throughout the course of a social exchange (Apicella et al., 2013; Carpenter, 2009; Stern, 2009). Synchrony has been widely applied across various fields to describe the temporal relationship between events that leads to coordinated states among the individual components of a system (Feldman, 2007; Leclère et al., 2014). In developmental science, synchrony has been especially useful for understanding the attunement between caregiver and child and predicting the child’s developmental outcomes (Leclère et al., 2014). Since its introduction, the concept of synchrony has expanded beyond behavioral events to encompass the coordination of physiological rhythms (Feldman, 2012). More recently, emerging research in human development has extended the concept of biobehavioral synchrony to include the temporal alignment of neural signals, laying the groundwork for studies on interpersonal neural synchrony (Carollo and Esposito, 2024; Carollo et al., 2021).

The emergence of the second-person neuroscience approach and hyperscanning, especially with functional near-infrared spectroscopy (fNIRS), has significantly advanced research on bio-behavioral synchrony, providing strong evidence for the existence of interpersonal neural synchrony (Carollo and Esposito, 2024; Carollo et al., 2021; Hakim et al., 2023; Hasson et al., 2012). Beyond single-participant studies (e.g., Rizzolatti and Fabbri-Destro 2008; Van Overwalle and Baetens 2009), the systematic review and meta-analysis by Czeszumski et al. (2022) reported that human cooperation is supported by a consistent and robust pattern of interpersonal neural synchrony in both frontal and temporoparietal regions, with stronger effects in the prefrontal cortex. Importantly, these effects were observed across diverse cooperation tasks, underscoring that these regions play a central role in supporting cooperative behavior rather than reflecting task-specific activations (Czeszumski et al., 2022). While cooperative tasks appear to induce synchrony in a network of fronto-temporo-parietal regions (e.g., Czeszumski et al. 2022; Wang et al. 2025), Czeszumski et al. (2022) highlighted that the type of cooperative task modulates which regions are most prominently engaged in neural synchrony. For example, the inferior and middle frontal gyri exhibit statistically significant interpersonal neural synchrony during gamified tasks, such as cooperative Jenga (Li et al., 2021; Liu et al., 2016). For instance, in their pioneering work, Cui et al. (2012) demonstrated the benefits of using fNIRS in a hyperscanning paradigm to explore the neural dynamics underlying real-life social interactions. In their study, participants played a cooperative video game side-by-side while their brain activity was monitored via fNIRS. The results revealed that interpersonal neural synchrony in the right superior frontal cortex was higher during the cooperative task compared to the competitive or solo conditions. Furthermore, greater synchrony was associated with better cooperation performance. Following this, other studies using fNIRS hyperscanning have investigated interpersonal neural synchrony in caregiver-child dyads. For example, Azhari et al. (2021) found increased interpersonal neural synchrony in left prefrontal regions during the co-exposure to animation videos in father-child dyads as compared to surrogate dyads. Similarly, Nguyen et al. (2021b, 2020) reported higher interpersonal synchrony in the inferior frontal gyrus (IFG) and temporoparietal junction (TPJ) during cooperative tasks in mother-child and father-child dyads compared to the individual performance of the same task. Moreover, in mother-child dyads, higher synchrony during cooperative problem-solving was associated with greater behavioral reciprocity and task success. Additionally, Nguyen et al. (2023, 2021a) observed a relationship between the rhythm of early interactions and interpersonal neural synchrony in mother-child dyads. In particular, they observed that, during proto-conversations with young infants, frequent turn-taking was associated with higher levels of interpersonal neural synchrony, although the strength of this relationship diminished as the proto-conversation progressed. In contrast, during full verbal conversations with five-year-olds, higher neural synchrony was correlated with more frequent turn-taking, primarily toward the end of the interaction. Altogether, these findings emphasize interpersonal neural synchrony as a potential biomarker of and quantitative measure for mother-child interaction quality (Nguyen et al., 2020; Reindl et al., 2018). Consistent with this, the electroencephalography (EEG) study by Endevelt-Shapira and Feldman (2023) showed that, in the theta frequency band, greater maternal sensitivity was associated with increased interpersonal neural synchrony, whereas higher levels of maternal intrusiveness were linked to reduced synchrony. Notably, synchrony between the mother’s right frontal and the infant’s left temporal regions was particularly sensitive to maternal sensitivity, while synchrony between the mother’s left frontal and the infant’s right temporal regions was indicative of intrusive caregiving.

The literature on bio-behavioral synchrony is by no means limited to caregiver-child dyads. The concept of interpersonal neural synchrony has been applied across a variety of social relationships, including romantic partners, close friends, colleagues, and even strangers (Astolfi et al., 2011; Carollo et al., 2025; Feldman, 2017; Gvirts and Perlmutter, 2020; Lim et al., 2024a; Long et al., 2021; Toppi et al., 2016). Moreover, hyperscanning studies have employed a wide array of tasks to explore interpersonal neural synchrony across diverse social contexts. Synchrony has been investigated in contexts where individuals co-attend to the same stimulus (e.g., watching a video together; Azhari et al., 2019; De Felice et al., 2024b), collaborate on cooperative tasks (e.g., Cui et al., 2012; Liu et al., 2021), or engage in conversational exchanges (e.g., Carollo et al., 2025; Nguyen et al., 2021a). These tasks vary in the degree of interactivity they require from participants. For instance, experimental conditions that involve co-attending to the same sensory stimulus demand minimal active social interaction, as participants are involved in a shared social context without engaging in direct exchanges. In contrast, tasks such as solving cooperative problems or participating in conversations require higher levels of interactivity and active social coordination. Furthermore, highly interactive scenarios can range from constrained, rules-based exchanges to completely free and naturalistic social interactions.

Consistently, De Felice et al. (2024a) identified two key dimensions central to the study of interpersonal neural synchrony in the literature: interpersonal closeness and interactivity. Interpersonal closeness refers to the degree of social proximity between interacting agents, often assessed through measures of relationship duration and/or intimacy. Interactivity, on the other hand, captures the level of reciprocity and the richness of multisensory mutual exchanges during a social interaction. Although both interpersonal closeness and interactivity are thought to enhance interpersonal neural synchrony, few studies have directly examined how these factors differentially contribute to synchrony across varying dyads and social contexts.

Nevertheless, some studies have explored how interpersonal neural synchrony varies across different dyad types, such as pairs of strangers, romantic partners, and caregiver-child. For example, using fNIRS, Pan et al. (2017) found that romantic partners exhibited stronger neural synchrony in the right superior frontal cortex compared to both friend and stranger dyads. Similarly, Song et al. (2024) reported higher synchrony between friends than between strangers during a cooperative task, particularly in the dorsolateral and medial regions of the prefrontal cortex. In a receiver-sender game paradigm, Shao et al. (2023) similarly found that romantic couples showed higher interpersonal neural synchrony in the frontopolar cortex and the right TPJ compared to stranger dyads. Initial comparisons between different relationship types have also been investigated using EEG. For instance, Kinreich et al. (2017) found significant neural synchrony in romantic partners, but not in strangers, localized in temporo-parietal areas and expressed in gamma-band activity. In a related study, Djalovski et al. (2021) observed a gradient of synchrony intensity across frequency bands (i.e., alpha, beta, gamma), with the strongest synchrony observed in romantic partners, followed by friends, and then strangers. At the physiological level, Bizzego et al. (2019a) found an opposite pattern, reporting a negative association between interpersonal closeness and physiological synchrony when comparing pairs of strangers, friends, and romantic partners during a co-watching task. While these studies provide preliminary evidence for the relationship between interpersonal closeness and neural synchrony, they typically focus on specific dyad comparisons and often do not vary task type.

Other studies have instead focused on how task demands shape interpersonal synchrony. For instance, Fishburn et al. (2018) used fNIRS and compared synchrony during passive co-watching and active cooperation, demonstrating that interactive tasks elicit higher levels of neural synchrony in the inferior and middle frontal gyri than passive ones. Likewise, Cui et al. (2012) and Liu et al. (2021) showed increased synchrony during joint problemsolving compared to control conditions involving shared attention without coordination. These findings suggest that the degree of required interactivity substantially modulates synchrony, likely due to increased reciprocal engagement and mutual prediction during real-time social exchanges.

Despite the insights gained from previous research, a systematic exploration of how interpersonal closeness and interactivity together shape and influence interpersonal neural synchrony remains an open question. This gap limits the cross-comparability of hyperscanning studies and, ultimately, poses challenges for developing a more integrative understanding of interpersonal synchrony. Notably, in their EEG study, Djalovski et al. (2021) compared different dyad types (i.e., romantic couples, best friends, and strangers) across both a motor and an empathy-based task, highlighting the value of such comparative designs. Building on this prior work, the present study extends the comparative approach by examining how interpersonal neural synchrony varies according to *(i)* the level of interpersonal closeness and *(ii)* the degree of social interactivity required by the experimental task. To achieve this, we employed fNIRS hyperscanning to monitor brain activity in three types of dyads: close friends, romantic partners, and mother-child pairs. These dyads were selected because they represent the most central and theoretically grounded relationships in the literature on interpersonal synchrony across multiple levels of analysis, as well as in developmental research. These dyads participated in three tasks designed to vary in interactivity: video coexposure, a rules-based cooperative game, and free verbal interaction. By varying both interpersonal closeness and task interactivity, this study aims to investigate how these factors shape interpersonal neural synchrony and to provide evidence that may inform future investigations into the neural mechanisms of human social interactions. Based on the bio-behavioral synchrony literature (De Felice et al., 2024a; Feldman, 2017), we hypothesized that:

- Interpersonal neural synchrony would be higher in dyads with greater interpersonal closeness, with the order being mother-child pairs having the highest synchrony, followed by romantic partners, and then close friends.
- Interpersonal neural synchrony would be influenced by the level of interactivity in the task, with lower synchrony observed in passive tasks (video co-exposure), higher synchrony in constrained social interactions (rules-based cooperative game), and the highest synchrony during more free-form interactions (free verbal interaction).

## 2. Methods

### 2.1. Study Design

All data for this study were collected using a hyperscanning approach, where dyads of participants took part in the experimental tasks while their brain activity was simultaneously monitored using fNIRS.

To examine the impact of interpersonal closeness on interpersonal neural synchrony, the study included three types of dyads: close friends, romantic partners, and mother-child pairs. These relationship types were chosen because they are the most consistently investigated in the literature on interpersonal synchrony and developmental psychology, are supported by a strong theoretical framework, and represent relationships that are central in individual development. Data collection for close friends and romantic partners took place in Italy at the University of Trento, while data for mother-child dyads was collected at the University of Vienna in Austria.

In the experiment, dyads of participants completed three distinct social tasks: video co-exposure, rules-based cooperative game, and free verbal interaction. These tasks were designed to vary the levels of interactivity, ranging from structured, passive activities to spontaneous, active exchanges.

The study was approved by the Ethics Committees of the University of Trento (ref. 2022-052) and the University of Vienna (ref. 00732). The experiment was conducted following the guidelines provided by the Declaration of Helsinki. Informed consent was obtained from all participants, and for the children involved, consent was provided by their primary caregivers.

### 2.2. Participants

A total of 284 participants (142 dyads) took part in the study. Specifically, the sample consisted of 70 dyads of close friends (29 male-male dyads, 23 female-female dyads, and 18 male-female dyads; Mean age = 21.60 ± 2.06 years), 39 heterosexual romantic partner dyads (Mean age = 23.45 ± 3.08 years), and 33 mother-child dyads (20 female children, Mean age mother = 38.59 ± 4.66 years, Mean age children = 5 years 6 months 18 days ± 3 months 22 days). A total of two close friend dyads, one romantic partner dyad, and two mother-child dyads were excluded for technical issues during the experiment.

To participate in the data collection as close friends, each dyad needed to have a friendship with no current or past romantic relationship between the two individuals. For romantic partner dyads, only heterosexual couples with an ongoing romantic relationship were included in the study. Finally, for the mother-child dyads, only mothers of children aged 5 years old were eligible for enrollment in the study. All participants reported no history of known or diagnosed health or neurological conditions, particularly those that could affect the oxygen-binding capacity of the blood.

Close friend and romantic partner dyads were recruited through convenience and snowball sampling via social media platforms and the University of Trento’s participant management and recruitment system (SONA). Mother-child dyads were recruited from a pre-existing database of volunteers at the University of Vienna. While close friend and romantic partner dyads did not receive monetary compensation, mother-child dyads were compensated with €6 (equivalent to the cost of two round-trip journeys on public transportation in Vienna) and a small toy for the child (i.e., a €6 Lego set).

### 2.3. Experimental Tasks

For all dyads, the experiment began with a resting-state measurement (2 minutes for adult dyads and 1 minute for mother-child dyads), which served as a baseline neural activity assessment. Additionally, mother-child dyads had a 1-minute resting-state measurement between each task. The distance between participants was approximately 60–80 cm for all dyads, and members of each dyad were always seated in separate chairs. This setup was consistent across conditions and dyad types.

The experiment consisted of three experimental tasks: video co-exposure, rules-based cooperative game, and free verbal interaction. These conditions were designed to explore the effect of interactivity levels on interpersonal neural synchrony, ranging from highly structured (video co-exposure) to semistructured (rules-based cooperative game) to unstructured (free verbal interaction) social interactions.

In the video co-exposure condition, a 3-minute, language-free animated video was presented to the dyads on a monitor positioned in front of them. The video was carefully selected to ensure consistency across experiments conducted in two different countries, thereby avoiding potential confounding variables related to language. It was also chosen for its appropriateness for both adult and child participants, making it suitable for all dyadic groups involved in the study. During this condition, dyads were instructed to simply attend to the video without interacting with each other. This setup allowed for the measurement of interpersonal neural synchrony driven by the shared sensory experience of the video in a social setting, without the influence of direct social interaction.

The rules-based cooperative condition involved a 5-minute session of cooperative Jenga. A Jenga tower was constructed and placed on a table between the participants, who were instructed to play according to the provided rules. Dyads were asked to take turns removing one block at a time and placing it on top of the tower. To maintain consistent difficulty across all dyads, participants were instructed to remove only one block per layer, starting from the bottom of the tower. In cases where the tower collapsed, participants were allowed to rebuild it and continue playing to ensure that the total duration of the task remained consistent for all dyads. This rule-based task was included to provide a semi-structured social interaction, ensuring a consistent level of predictability of the social exchange.

In the free verbal interaction condition, participants sat facing each other and were free to engage in verbal interaction as they preferred for a duration of 5 minutes. This unstructured task allowed for natural, spontaneous communication between the dyads. Whereas cooperative Jenga constrains interaction within fixed rules and predictable turn-taking mainly oriented toward the game, free verbal interaction allows for open-ended and less predictable exchanges oriented toward mutual communication.

The order of the experimental conditions was randomized for each dyad to mitigate potential order effects on the results. For the current work, we did not analyze the resting-state data.

### 2.4. Acquisition of Neural Data

fNIRS hyperscanning was used across all experimental tasks to monitor participants’ brain activity. Each participant wore a cap equipped with eight LED sources emitting light at wavelengths of 760 nm and 850 nm, along with eight detectors arranged to capture brain activity from the bilateral IFG and bilateral TPJ, following previous research protocols (e.g., Lei et al., 2025; Nguyen et al., 2020).

The cap configurations differed slightly across data collections. In the close friends and romantic partners studies conducted in Italy, one detector was dedicated to collecting signals from eight short-distance channels, resulting in a total of 14 fNIRS channels per participant. In contrast, for the mother-child dyads study conducted in Vienna, short-distance channels were not available. Instead, the eighth detector was used to add two long-distance channels, leading to a total of 16 fNIRS channels per participant.

For the data analysis, fNIRS channels were aggregated into four regions of interest: left IFG, right IFG, left TPJ, and right TPJ. The cap setup for the close friends and romantic partners studies was optimized using the fNIRS Optodes’ Location Decider (fOLD) to maximize sensitivity in the target regions of interest (Zimeo Morais et al., 2018) (see Figure S1A in the Supplementary Materials). Due to logistical constraints in the laboratory in Austria, the cap setup for the mother-child dyads study was designed based on previous adult-child studies (Nguyen et al., 2021a,b, 2020) (see Figure S1B in the Supplementary Materials).

For the experiment, channel placement followed the international 10-20 EEG system. In the data collection conducted in Italy, optode stabilizers ensured that the distance between sources and detectors never exceeded 3 cm, facilitating the acquisition of data with a high signal-to-noise ratio. Additionally, different fNIRS devices were used across locations: a NIRSport2 device (NIRx Medical Technologies LLC) with a sampling rate of 10.17 Hz was used in Italy, while a NIRSport device (NIRx Medical Technologies LLC) with a sampling rate of 7.81 Hz was used in Austria.

### 2.5. Processing of Neural Data

Pre-processing and synchrony analyses were conducted using *pyphysio* (Bizzego et al., 2019b). The fNIRS raw intensity signals were first converted to changes in optical density. To ensure consistency in the length of data across experimental conditions, the optical density data were segmented to include only the first 3 minutes of each condition. Signal quality was assessed using a convolutional neural network model trained to classify the quality of fNIRS segments (Bizzego et al., 2022, 2021). The processed data were then resampled to 10 Hz and converted into oxygenated (HbO) and deoxygenated hemoglobin (HbR) values using the modified Beer-Lambert law (Delpy et al., 1988). For the statistical analyses, we focused exclusively on HbO values, consistent with previous studies (e.g., Lim et al., 2024a,b; see the results on HbR values in the Supplementary Materials). Finally, temporal autocorrelations and physiological noise in the neural data were removed by whitening the signals through an autoregressive model (Barker et al., 2013). The use of an autoregressive filter to pre-whiten the signals before the analysis has been shown to reduce the false discovery rate when estimating the coherence between signals (Santosa et al., 2017).

### 2.6. Interpersonal Neural Synchrony

Interpersonal neural synchrony was calculated on a channel-by-channel basis between participants using wavelet transform coherence (WTC) (Grinsted et al., 2004). WTC allows estimating the coherence between fNIRS time series considering both frequency and time dimensions, while also accounting for both in-phase and phase-lagged correlations (Nguyen et al., 2020). In this way, WTC facilitates a comprehensive analysis of global coherence patterns in brain activity.

For each pair of good-quality channels in every experimental task, WTC was calculated over a frequency range of 0.01 to 0.20 Hz, following the approach of previous works (Carollo et al., 2025; Lim et al., 2024a,b). To obtain the final WTC value, coherence scores were averaged across the entire frequency spectrum. This approach ensured an unbiased analysis, as no specific frequency bands were hypothesized *a priori* (Carollo et al., 2025).

To obtain a baseline level of synchrony between participants who did not interact directly with one another but did the same experimental task, we also computed WTC between non-interacting surrogate dyads. These were constructed by randomly pairing participants from different original dyads within the same dataset and dyad type (e.g., friend with friend, mother with child), ensuring no cross-cohort pairings occurred. As in Carollo et al. (2025), we performed a single permutation of the dyads, resulting in the same number of data points as the true dyads. To maintain consistency in experimental conditions and fNIRS channel alignment, we formed each surrogate pair by matching a participant’s data segment from a given channel and task (e.g., channel X during video co-exposure) with the corresponding segment from another participant within the same condition. For example, if participant A from dyad 001 had fNIRS data from channel X during the video co-exposure task, we paired it with the fNIRS data from channel X of participant B from dyad 002 recorded during the same task. Synchrony observed in surrogate dyads is assumed to reflect stimulus-driven effects alone, allowing us to separate them from the additional synchrony due to co-presence in true dyads (Golland et al., 2015).

### 2.7. Data Analysis

The data analysis for the current study is organized into three main steps:

1. Identifying the combinations of brain regions between the two participants where a statistically significant difference in WTC scores exists between true and surrogate dyads;
2. Outline the effect of interpersonal closeness and social interactivity on overall interpersonal neural synchrony across all combinations of regions of interest between the two participants;
3. Outline the effect of interpersonal closeness and social interactivity on interpersonal neural synchrony in specific combinations of brain regions between the two participants.

Across analytical steps, we used linear mixed models (Bates et al., 2005), which are particularly well suited for unbalanced designs as they can appropriately handle unequal group sizes and missing data structures. Before conducting the analyses, we created two types of region combination IDs: one that identified combinations of brain regions regardless of their order (e.g., treating region X in participant A–region Y in participant B and region Y in participant A–region X in participant B as equivalent) and another that accounted for the specific order of the combination (e.g., treating region X in participant A–region Y in participant B and region Y in participant A–region X in participant B as distinct). Subsequently, we conducted:

1. A separate linear mixed model for each combination of regions of interest (independently from their order) between the two participants, with WTC scores as the dependent variable and dyad type (true *vs*. surrogate dyads) as the fixed effect. Dyadic relationships (i.e., close friends, romantic partners, mother-child dyads), experimental conditions (i.e., video co-exposure, rules-based cooperative game, free verbal interaction), and dyad IDs were included as random effects. For models based on synchrony data from combinations of non-homologous brain regions, we also included the order-dependent region combination IDs as a random effect. A Bonferroni correction, chosen for its conservative control of false positives in the presence of multiple comparisons, was applied.
2. A single linear mixed model using data from all brain region combinations. This model was designed to examine the overall effect of interpersonal closeness and social interactivity on interpersonal neural synchrony, using only data from true dyads. WTC scores in true dyads served as the dependent variable, while dyadic relationships and experimental tasks were included as fixed effects. The model also included the two types of region combination IDs (one that is order-independent and one that is order-dependent) and dyad IDs as random effects. *Posthoc* pairwise comparisons of marginal means were performed to identify significant differences between levels of each significant main factor. A Bonferroni correction was applied to control the false positive rate in *post-hoc* comparisons.
3. A separate linear mixed model with data from each combination of regions of interest (independently from their order) between the two participants. In these models, WTC scores from true dyads were the dependent variable, and dyadic relationships and experimental conditions were used as fixed effects. Additionally, the model included the dyad IDs and, only for models conducted on non-homologous combinations of brain region, the order-dependent region combination IDs as random effects. *Post-hoc* pairwise comparisons of marginal means were performed, with Bonferroni correction applied to control for multiple comparisons and minimize the risk of Type I error.

## 3. Results

### 3.1. Neural Synchrony in True Versus Surrogate Dyads

In the first stage of the analysis, we used a linear mixed model to test the differences in interpersonal neural synchrony between true and surrogate dyads for each combination of brain regions between the two participants. The results revealed significant differences in synchrony across all brain region combinations, with true dyads showing higher levels of synchrony compared to surrogate dyads (i.e., left IFG–left IFG: *t* = 12.48, *p* <.001, *q* <.001, *d* = 0.24, 95% CI [0.20, 0.27]; left IFG–right IFG: *t* = 18.60, *p* <.001, *q* <.001, *d* = 0.25, 95% CI [0.22, 0.28]; left IFG–left TPJ: *t* = 16.48, *p* <.001, *q* <.001, *d* = 0.22, 95% CI [0.20, 0.25]; left IFG–right TPJ: *t* = 13.15, *p* <.001, *q* <.001, *d* = 0.23, 95% CI [0.19, 0.26]; right IFG–right IFG: *t* = 16.81, *p* <.001, *q* <.001, *d* = 0.32, 95% CI [0.29, 0.36]; right IFG–left TPJ: *t* = 19.20, *p* <.001, *q* <.001, *d* = 0.26, 95% CI [0.23, 0.29]; right IFG–right TPJ: *t* = 13.77, *p* <.001, *q* <.001, *d* = 0.24, 95% CI [0.21, 0.28]; left TPJ–left TPJ: *t* = 11.80, *p* <.001, *q* <.001, *d* = 0.23, 95% CI [0.19, 0.27]; left TPJ–right TPJ: *t* = 10.20, *p* <.001, *q* <.001, *d* = 0.18, 95% CI [0.15, 0.21]; right TPJ–right TPJ: *t* = 5.42, *p* <.001, *q* <.001, *d* = 0.17, 95% CI [0.11, 0.23]; see Figure 1). This finding suggests that the neural synchrony observed in true dyads is distinct from surrogate, noninteracting dyads, especially when looking at the right IFG, supporting the hypothesis that dyadic interactions contribute to meaningful interpersonal neural synchrony. Given these statistical differences, we used all data from true dyads across all combinations of brain regions for subsequent analysis.

**Figure 1.**
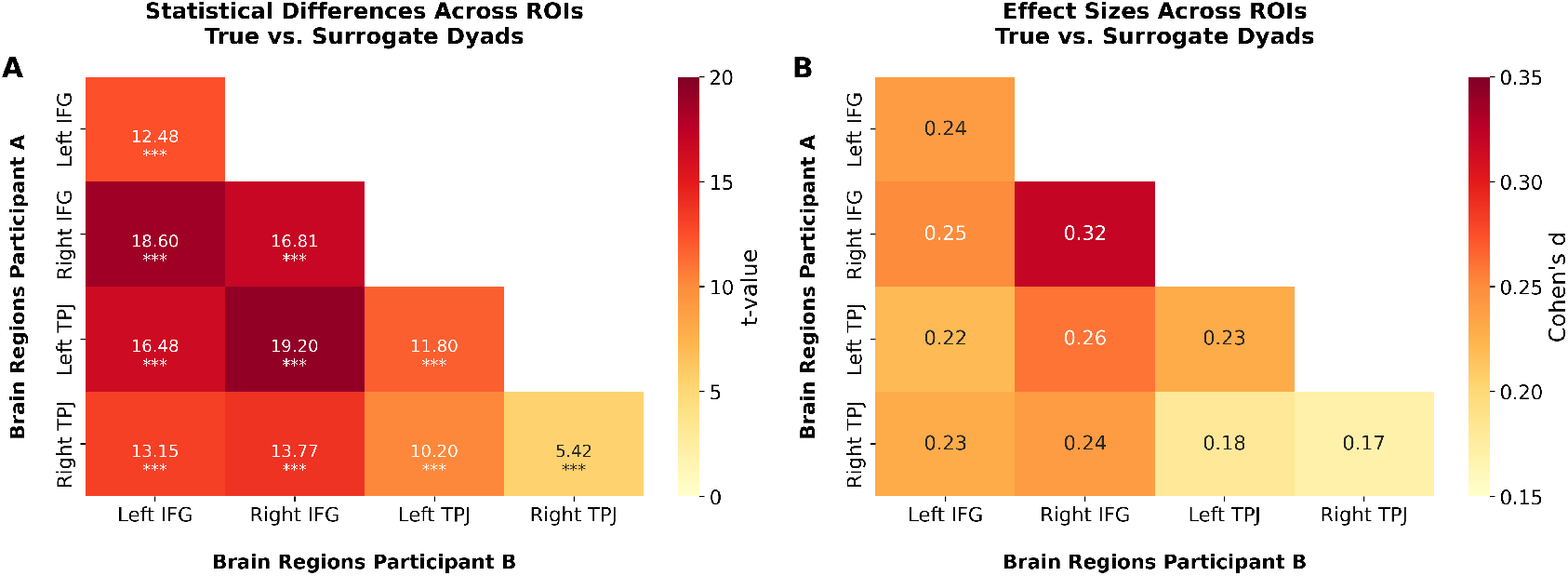
**(A)** Statistical differences between true and surrogate dyads in terms of interpersonal neural synchrony. For each pairwise combination of brain regions of interest (ROIs; i.e., bilateral inferior frontal gyri (IFG) and bilateral temporoparietal junctions (TPJ)) between the two participants involved in the experiment, we report the *t* -value and its significance level. A linear mixed model was used with wavelet transform coherence scores as the dependent variable and dyad type (true *vs*. surrogate) as the fixed effect. Dyadic relationships, experimental conditions, and dyad IDs were included as random effects. For non-homologous region combinations, an order-dependent region combination ID was also included as a random effect. Bonferroni correction was applied to adjust the alpha level for multiple comparisons (^***^ *q* <.001). **(B)** Effect sizes (Cohen’s *d*) corresponding to the statistical contrasts shown in panel **A**.

### 3.2. Interpersonal Closeness and Social Interactivity on Neural Synchrony (All Combinations of Regions)

In the second stage of our analysis, we conducted a linear mixed model to examine the effects of interpersonal closeness and levels of social interactivity on interpersonal neural synchrony across all regions of interest. Our results indicate that both interpersonal closeness (*F* (2, 131) = 10.60, *p* <.001, partial *η*^2^ = 0.14, 95% CI [0.05, 1.00]; see Figure 2A) and social interactivity (*F* (2, 71,653) = 185.81, *p* <.001, partial *η*^2^ = 0.01, 95% CI [0.00, 1.00]; see Figure 2B) have a statistically significant effect on interpersonal neural synchrony scores. Additionally, a statistically significant interaction effect between interpersonal closeness and social interactivity on interpersonal neural synchrony emerged (*F* (4, 71,654) = 69.14, *p* <.001, partial *η*^2^ = 0.01, 95% CI [0.00, 1.00]).

**Figure 2.**
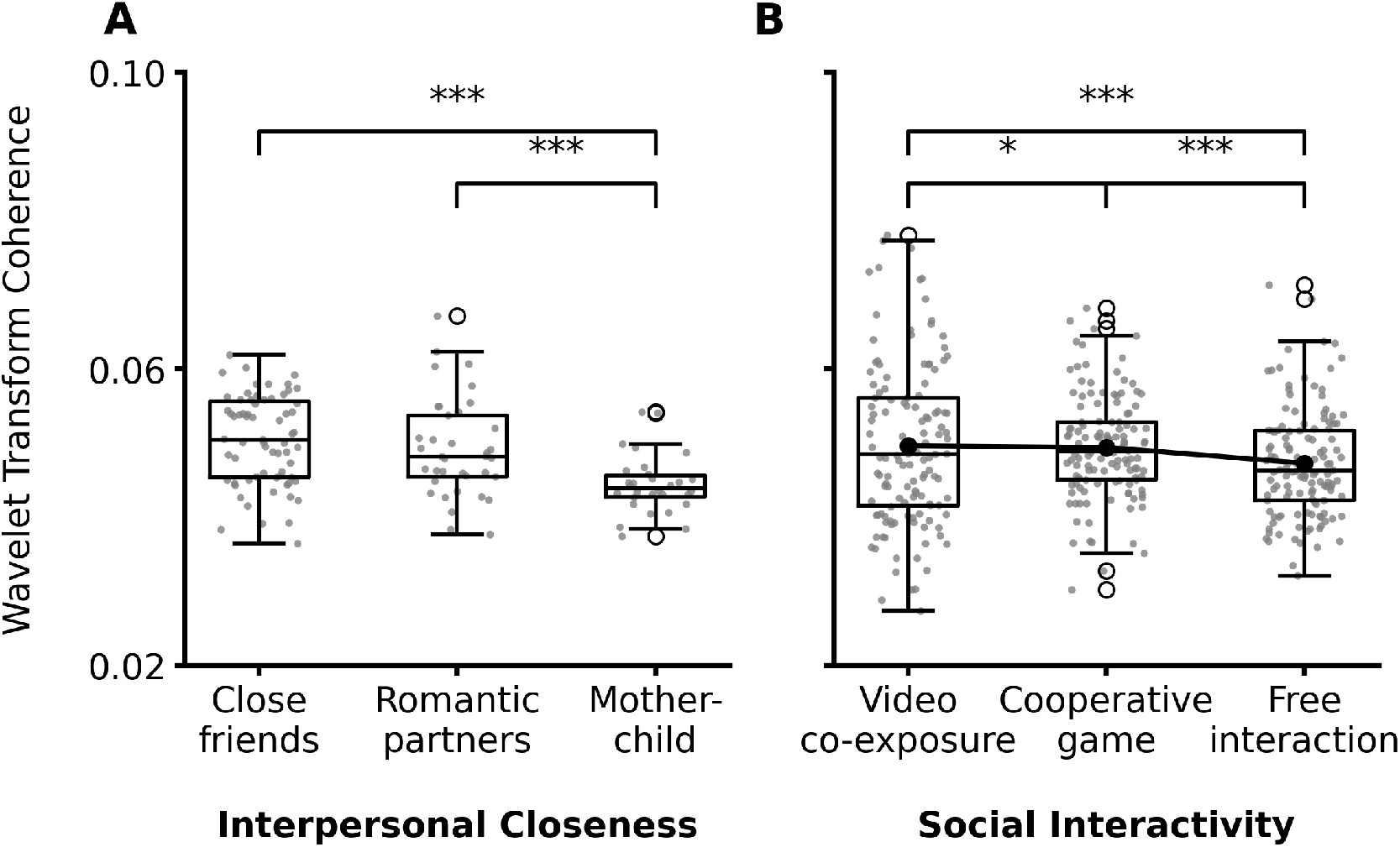
**(A)** Comparison of overall interpersonal neural synchrony scores across levels of interpersonal closeness (i.e., close friends, romantic partners, and mother-child dyads). **(B)** Comparison of overall interpersonal neural synchrony scores across levels of social interactivity (i.e., video co-exposure, rules-based cooperative game, and free verbal interaction). (^*^ *q<*.05; ^***^ *q* <.001).

Pairwise comparisons of marginal means revealed significant differences in interpersonal neural synchrony between close friend dyads and mother-child dyads (estimate = 0.00586, Standard Error (SE) = 0.00128, *z* = 4.57, *q* <.001, *d* = 0.42, 95% CI [0.24, 0.60]), as well as between romantic partners and mother-child dyads (estimate = 0.00466, SE = 0.00143, *z* = 3.26, *q* =.003, *d* = 0.33, 95% CI [0.13, 0.53]). However, no significant difference was found between close friend dyads and romantic partner dyads (*q >*.05).

Further *post-hoc* analyses showed significant differences in interpersonal neural synchrony between video co-exposure and rules-based cooperative game (estimate = 0.000353, SE = 0.000136, *z* = 2.59, *q* =.029, *d* = 0.03, 95% CI [0.01, 0.04]), video co-exposure and free verbal interaction (estimate = 0.002411, SE = 0.000135, *z* = 17.82, *q* <.001, *d* = 0.17, 95% CI [0.15, 0.19]), and rules-based cooperative game and free verbal interaction (estimate = 0.002058, SE = 0.000136, *z* = 15.18, *q* <.001, *d* = 0.15, 95% CI [0.13, 0.17]).

*Post-hoc* analyses of the interaction between interpersonal closeness and social interactivity on interpersonal neural synchrony revealed distinct patterns across dyad types and tasks. When examining task-related differences within each dyad type, close friends and romantic partners showed a consistent trend: synchrony was highest during video co-exposure, followed by the cooperative game, and lowest during free interaction, in line with the pattern observed in the main effects. In contrast, mother–child dyads displayed no statistically significant difference in interpersonal neural synchrony across levels of social interactivity. Table 1 reports the complete set of comparisons and statistics for these effects.

**Table 1:** Results from the *post-hoc* pairwise comparisons of the interaction between interpersonal closeness and social interactivity on interpersonal neural synchrony (across all brain region combinations). For each level of one factor, the table reports the estimated contrasts between two levels of the other factor, together with the standard error, *z* -ratio, corrected *q* -value, and Cohen’s *d*. (^*^ *q* <.05; ^**^ *q* <.01; ^***^ *q* <.001).

| Contrast | Estimate | Standard Error | z-ratio | q | Cohen's *d* 95% CI |
| --- | --- | --- | --- | --- | --- |
| **Close friends** |  |  |  |  |  |
| Video-Cooperative game | 0.00061 | 0.0002 | 3.28 | .038 * | 0.04 [0.02, 0.07] |
| Video-Free interaction | 0.00216 | 0.0002 | 11.94 | < .001 *** | 0.15 [0.13, 0.18] |
| Cooperative game-Free interaction | 0.00155 | 0.0002 | 8.35 | < .001 *** | 0.11 [0.08, 0.14] |
| **Romantic partners** |  |  |  |  |  |
| Video-Cooperative game | 0.00090 | 0.0003 | 3.60 | .011 * | 0.06 [0.03, 0.10] |
| Video-Free interaction | 0.00537 | 0.0002 | 21.53 | < .001 *** | 0.38 [0.35, 0.42] |
| Cooperative game-Free interaction | 0.00447 | 0.0003 | 17.68 | < .001 *** | 0.32 [0.28, 0.35] |
| **Mother-child** |  |  |  |  |  |
| Video-Cooperative game | -0.00045 | 0.0003 | -1.71 | > .05 | / |
| Video-Free interaction | -0.00030 | 0.0003 | -1.12 | > .05 | / |
| Cooperative game-Free interaction | 0.00016 | 0.0003 | 0.61 | > .05 | / |
| **Video co-exposure** |  |  |  |  |  |
| Close friends-Romantic partners | 0.00003 | 0.0012 | 0.02 | > .05 | / |
| Close friends-Mother-child | 0.00703 | 0.0013 | 5.43 | < .001 *** | 0.50 [0.32, 0.68] |
| Romantic partners-Mother-child | 0.00700 | 0.0014 | 4.84 | < .001 *** | 0.50 [0.30, 0.70] |
| **Cooperative game** |  |  |  |  |  |
| Close friends-Romantic partners | 0.00032 | 0.0012 | 0.26 | > .05 | / |
| Close friends-Mother-child | 0.00597 | 0.0013 | 4.61 | < .001 *** | 0.43 [0.25, 0.61] |
| Romantic partners-Mother-child | 0.00565 | 0.0014 | 3.90 | .004 ** | 0.40 [0.20, 0.61] |
| **Free interaction** |  |  |  |  |  |
| Close friends-Romantic partners | 0.00324 | 0.0012 | 2.68 | > .05 | / |
| Close friends-Mother-child | 0.00458 | 0.0013 | 3.54 | .015 * | 0.33 [0.15, 0.51] |
| Romantic partners-Mother-child | 0.00134 | 0.0014 | 0.92 | > .05 | / |

Additionally, in both the video co-exposure and cooperative game conditions, mother–child dyads showed the lowest levels of synchrony compared to close friends and romantic partners, with no statistically significant differences between the latter two groups. In the free interaction condition, only close friends exhibited higher synchrony than mother–child dyads, while no statistically significant differences emerged between close friends and romantic partners, nor between romantic partners and mother–child dyads. The full set of comparisons and statistics for these effects is reported in Table 1.

These findings suggest that both the nature of the relationship between individuals and the type of social interaction they engage in play crucial roles in modulating interpersonal neural synchrony. However, an inspection of the effect sizes indicates that interpersonal closeness may exert a stronger influence on interpersonal neural synchrony than the level of social interactivity.

### 3.3. Interpersonal Closeness and Social Interactivity on Neural Synchrony (Individual Combinations of Regions)

In the third stage of our analysis, we conducted a linear mixed model to examine the effects of interpersonal closeness and social interactivity on interpersonal neural synchrony across individual combinations of regions of interest. Our results indicate that both interpersonal closeness and social interactivity had a statistically significant effect on synchrony scores in the following brain region pairs: left IFG–left IFG, left IFG–right IFG, left IFG– left TPJ, right IFG–right IFG, right IFG–left TPJ, and right IFG–right TPJ.

Conversely, social interactivity—but not interpersonal closeness—had a statistically significant effect on synchrony in the following brain region pairs: left IFG–right TPJ, left TPJ–left TPJ, left TPJ–right TPJ, and right TPJ– right TPJ. Table 2 reports the results of the analysis investigating the effect of interpersonal closeness and social interactivity levels on interpersonal neural synchrony across combinations of regions of interest.

**Table 2:** Results of the analysis examining the main effects of interpersonal closeness and social interactivity on interpersonal neural synchrony across different combinations of brain regions. For each combination, we report the Sum of Squares (Sum Sq), Mean Square (Mean Sq), numerator and denominator degrees of freedom (NumDf, DenDf), *F* - value, *p*-value, corrected *q* -value, and partial *η*^2^. (^*^ *q* <.05; ^**^ *q* <.01; ^***^ *q* <.001).

| Factor | Sum Sq | Mean Sq | NumDf | DenDf | *F* | *p* | *q* | partial $\eta^2$ 95% CI |
| --- | --- | --- | --- | --- | --- | --- | --- | --- |
| **Left inferior frontal gyrus – left inferior frontal gyrus** |  |  |  |  |  |  |  |  |
| Interpersonal closeness | 0.003 | 0.001 | 2 | 130.2 | 8.31 | < .001 | .008 ** | 0.11 [0.04, 1.00] |
| Social interactivity | 0.013 | 0.006 | 2 | 5581.1 | 37.42 | < .001 | < .001 *** | 0.01 [0.01, 1.00] |
| **Left inferior frontal gyrus – right inferior frontal gyrus** |  |  |  |  |  |  |  |  |
| Interpersonal closeness | 0.008 | 0.004 | 2 | 132.4 | 20.98 | < .001 | < .001 *** | 0.24 [0.14, 1.00] |
| Social interactivity | 0.020 | 0.010 | 2 | 11,072.3 | 51.95 | < .001 | < .001 *** | 0.01 [0.01, 1.00] |
| **Left inferior frontal gyrus – left temporoparietal junction** |  |  |  |  |  |  |  |  |
| Interpersonal closeness | 0.003 | 0.001 | 2 | 131.8 | 7.54 | < .001 | .016 * | 0.10 [0.03, 1.00] |
| Social interactivity | 0.008 | 0.004 | 2 | 11,012.8 | 22.45 | < .001 | < .001 *** | 0.004 [0.00, 1.00] |
| **Left inferior frontal gyrus – right temporoparietal junction** |  |  |  |  |  |  |  |  |
| Interpersonal closeness | 0.002 | 0.001 | 2 | 128.3 | 4.86 | .009 | .185 | / |
| Social interactivity | 0.010 | 0.005 | 2 | 6603.0 | 29.80 | < .001 | < .001 *** | 0.01 [0.01, 1.00] |
| **Right inferior frontal gyrus – right inferior frontal gyrus** |  |  |  |  |  |  |  |  |
| Interpersonal closeness | 0.008 | 0.004 | 2 | 131.2 | 20.44 | < .001 | < .001 *** | 0.24 [0.13, 1.00] |
| Social interactivity | 0.021 | 0.011 | 2 | 5390.5 | 55.16 | < .001 | < .001 *** | 0.02 [0.01, 1.00] |
| **Right inferior frontal gyrus – left temporoparietal junction** |  |  |  |  |  |  |  |  |
| Interpersonal closeness | 0.006 | 0.003 | 2 | 132.2 | 14.51 | < .001 | < .001 *** | 0.18 [0.09, 1.00] |
| Social interactivity | 0.027 | 0.014 | 2 | 10,838.9 | 66.97 | < .001 | < .001 *** | 0.01 [0.01, 1.00] |
| **Right inferior frontal gyrus – right temporoparietal junction** |  |  |  |  |  |  |  |  |
| Interpersonal closeness | 0.004 | 0.002 | 2 | 127.5 | 9.75 | < .001 | .002 ** | 0.13 [0.05, 1.00] |
| Social interactivity | 0.003 | 0.001 | 2 | 6451.6 | 7.66 | < .001 | .010 * | 0.002 [0.00, 1.00] |
| **Left temporoparietal junction – left temporoparietal junction** |  |  |  |  |  |  |  |  |
| Interpersonal closeness | 0.0003 | 0.0002 | 2 | 128.7 | 0.84 | .434 | .999 | / |
| Social interactivity | 0.013 | 0.007 | 2 | 5337.3 | 36.71 | < .001 | < .001 *** | 0.01 [0.00, 1.00] |
| **Left temporoparietal junction – right temporoparietal junction** |  |  |  |  |  |  |  |  |
| Interpersonal closeness | 0.0005 | 0.0002 | 2 | 128.8 | 1.11 | 0.33 | .999 | / |
| Social interactivity | 0.005 | 0.002 | 2 | 6435.1 | 13.50 | < .001 | < .001 *** | 0.004 [0.00, 1.00] |
| **Right temporoparietal junction – right temporoparietal junction** |  |  |  |  |  |  |  |  |
| Interpersonal closeness | 0.0001 | 0.0001 | 2 | 119.14 | 0.35 | .705 | .999 | / |
| Social interactivity | 0.004 | 0.002 | 2 | 2014.71 | 13.06 | < .001 | < .001 *** | 0.01 [0.00, 1.00] |

Pairwise comparisons of marginal means revealed that close friends exhibited significantly higher interpersonal neural synchrony than mother-child dyads in the left IFG–left IFG, left IFG–right IFG, left IFG–left TPJ, right IFG–right IFG, right IFG–left TPJ, and right IFG–right TPJ. Moreover, romantic partners showed significantly higher brain synchrony than mother-child dyads in the left IFG–right IFG, right IFG–right IFG, right IFG–left TPJ, and right IFG–right TPJ. The complete results of these analyses are presented in Table 3 and represented in Figure 3. These findings suggest that interpersonal neural synchrony is generally stronger among close friends and romantic partners compared to mother-child dyads, as found in the analysis presented in Section 3.2.

**Table 3:**
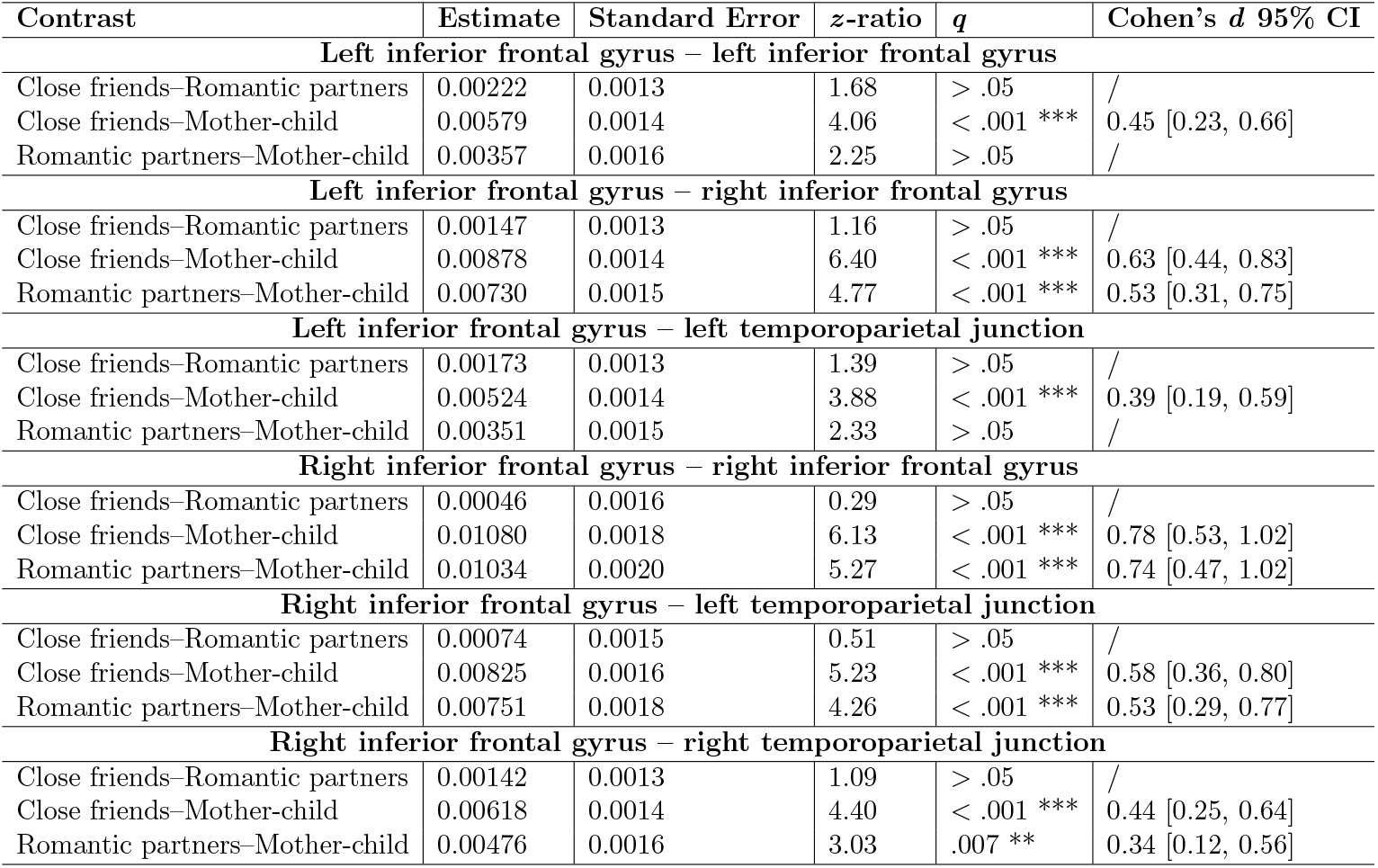
Results of the *post-hoc* pairwise comparisons examining the effect of interpersonal closeness on interpersonal neural synchrony across different brain region combinations. For each combination, we report the estimated contrast between two levels of interpersonal closeness, along with the standard error, *z* -ratio, corrected *q* -value, and Cohen’s *d*. (^**^ *q* <.01; ^***^ *q* <.001).

**Figure 3.**
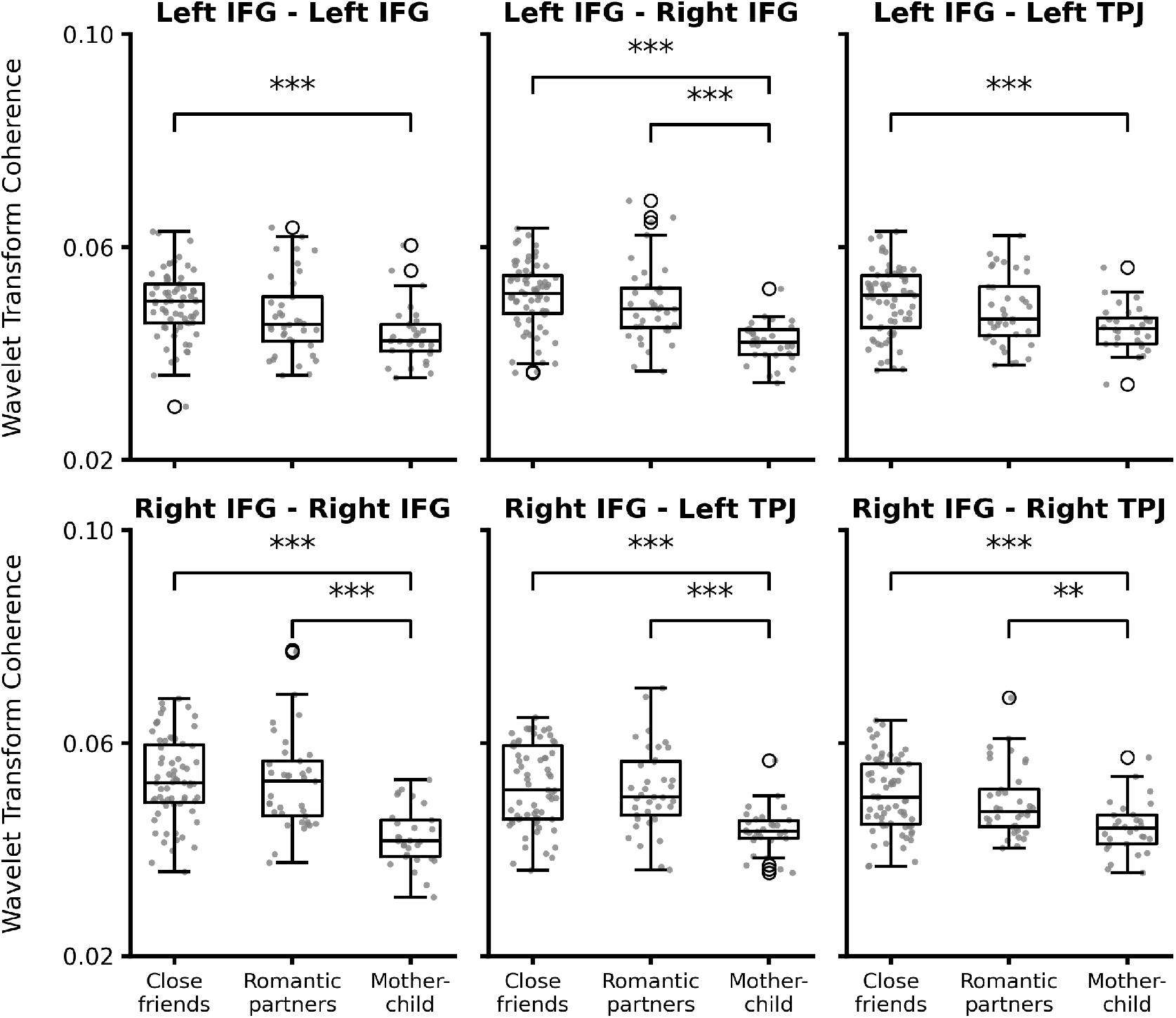
Effect of interpersonal closeness on interpersonal neural synchrony across combinations of regions of interest (i.e., bilateral inferior frontal gyri (IFG) and bilateral temporoparietal junctions (TPJ)). (^**^ *q* <.01; ^***^ *q* <.001).

Furthermore, regarding the effect of social interactivity, pairwise comparisons of marginal means revealed distinct synchrony patterns across different brain region pairs. Notably, the video co-exposure condition exhibited significantly greater synchrony than the other experimental tasks in the following combinations: right IFG–right IFG, right IFG–left TPJ, left TPJ–left TPJ, and left TPJ–right TPJ. In contrast, the rules-based cooperative game condition demonstrated the highest synchrony for left IFG–left IFG and left IFG–right TPJ. Additionally, both the video co-exposure and rules-based cooperative game conditions showed greater synchrony than free interaction in the following region pairs: left IFG–right IFG, left IFG–left TPJ, right IFG–right TPJ, and right TPJ–right TPJ. Overall, across all experimental tasks, free interaction consistently exhibited the lowest level of interpersonal neural synchrony. The full results detailing the effect of interactivity levels on neural synchrony are provided in Table 4 and represented in Figure 4.

**Table 4:** Results of the *post-hoc* pairwise comparisons examining the effect of social interactivity levels on interpersonal neural synchrony across different brain region combinations. For each combination, we report the estimated contrast between two levels of social interactivity, along with the standard error, *z* -ratio, and corrected *q* -value, and Cohen’s *d*. (^*^ *q* <.05; ^***^ *q* <.001).

| Contrast | Estimate | Standard Error | z-ratio | q | Cohen's d 95% CI |
| --- | --- | --- | --- | --- | --- |
| **Left inferior frontal gyrus – left inferior frontal gyrus** |  |  |  |  |  |
| Video-Cooperative game | -0.00203 | 0.0004 | -4.73 | < .001 *** | -0.16 [-0.22, -0.09] |
| Video-Free interaction | 0.00166 | 0.0004 | 3.91 | < .001 *** | 0.13 [0.06, 0.19] |
| Cooperative game-Free interaction | 0.00369 | 0.0004 | 8.64 | < .001 *** | 0.28 [0.22, 0.35] |
| **Left inferior frontal gyrus – right inferior frontal gyrus** |  |  |  |  |  |
| Video-Cooperative game | -0.000403 | 0.0003 | -1.23 | > .05 | / |
| Video-Free interaction | 0.002624 | 0.0003 | 8.15 | < .001 *** | 0.19 [0.14, 0.24] |
| Cooperative game-Free interaction | 0.003027 | 0.0003 | 9.32 | < .001 *** | 0.22 [0.17, 0.26] |
| **Left inferior frontal gyrus – left temporoparietal junction** |  |  |  |  |  |
| Video-Cooperative game | -0.000124 | 0.0003 | -0.40 | > .05 | / |
| Video-Free interaction | 0.001758 | 0.0003 | 5.61 | < .001 *** | 0.13 [0.09, 0.18] |
| Cooperative game-Free interaction | 0.001882 | 0.0003 | 5.94 | < .001 *** | 0.14 [0.09, 0.19] |
| **Left inferior frontal gyrus – right temporoparietal junction** |  |  |  |  |  |
| Video-Cooperative game | -0.00251 | 0.0004 | -6.25 | < .001 *** | -0.19 [-0.25, -0.13] |
| Video-Free interaction | 0.00032 | 0.0004 | 0.81 | > .05 | / |
| Cooperative game-Free interaction | 0.00284 | 0.0004 | 7.09 | < .001 *** | 0.21 [0.15, 0.27] |
| **Right inferior frontal gyrus – right inferior frontal gyrus** |  |  |  |  |  |
| Video-Cooperative game | 0.00220 | 0.0005 | 4.70 | < .001 *** | 0.16 [0.09, 0.22] |
| Video-Free interaction | 0.00484 | 0.0005 | 10.49 | < .001 *** | 0.35 [0.28, 0.41] |
| Cooperative game-Free interaction | 0.00264 | 0.0005 | 5.66 | < .001 *** | 0.19 [0.12, 0.26] |
| **Right inferior frontal gyrus – left temporoparietal junction** |  |  |  |  |  |
| Video-Cooperative game | 0.00253 | 0.0003 | 7.47 | < .001 *** | 0.18 [0.13, 0.23] |
| Video-Free interaction | 0.00379 | 0.0003 | 11.38 | < .001 *** | 0.27 [0.22, 0.31] |
| Cooperative game-Free interaction | 0.00127 | 0.0003 | 3.76 | < .001 *** | 0.09 [0.04, 0.14] |
| **Right inferior frontal gyrus – right temporoparietal junction** |  |  |  |  |  |
| Video-Cooperative game | -0.000583 | 0.0004 | -1.36 | > .05 | / |
| Video-Free interaction | 0.001057 | 0.0004 | 2.50 | .037 * | 0.08 [0.02, 0.14] |
| Cooperative game-Free interaction | 0.001640 | 0.0004 | 3.85 | < .001 *** | 0.12 [0.06, 0.18] |
| **Left temporoparietal junction – left temporoparietal junction** |  |  |  |  |  |
| Video-Cooperative game | 0.003630 | 0.0005 | 7.93 | < .001 *** | 0.27 [0.20, 0.33] |
| Video-Free interaction | 0.003039 | 0.0005 | 6.74 | < .001 *** | 0.23 [0.16, 0.29] |
| Cooperative game-Free interaction | -0.000591 | 0.0005 | -1.29 | > .05 | / |
| **Left temporoparietal junction – right temporoparietal junction** |  |  |  |  |  |
| Video-Cooperative game | 0.001644 | 0.0004 | 3.94 | < .001 *** | 0.12 [0.06, 0.18] |
| Video-Free interaction | 0.002009 | 0.0004 | 4.89 | < .001 *** | 0.15 [0.09, 0.21] |
| Cooperative game-Free interaction | 0.000365 | 0.0004 | 0.88 | > .05 | / |
| **Right temporoparietal junction – right temporoparietal junction** |  |  |  |  |  |
| Video-Cooperative game | -0.000603 | 0.0007 | -0.86 | > .05 | / |
| Video-Free interaction | 0.002724 | 0.0007 | 3.94 | < .001 *** | 0.21 [0.11, 0.31] |
| Cooperative game-Free interaction | 0.003327 | 0.0007 | 4.77 | < .001 *** | 0.26 [0.15, 0.36] |

**Figure 4.**
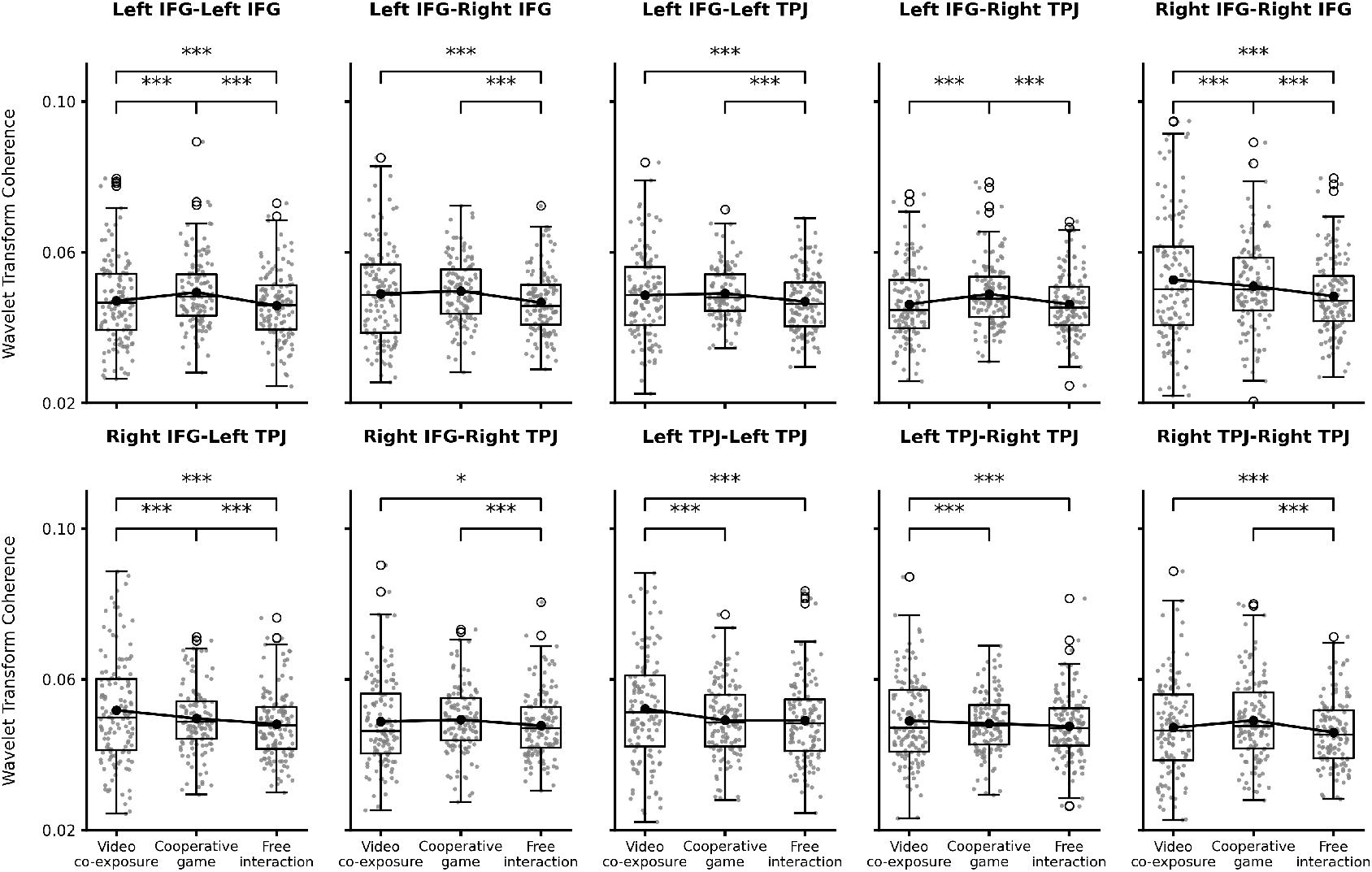
Effect of social interactivity on interpersonal neural synchrony across combinations of regions of interest (i.e., bilateral inferior frontal gyri (IFG) and bilateral temporoparietal junctions (TPJ)). (^*^ *q* <.05; ^***^ *q* <.001).

## 4. Discussion

In this study, we conducted an fNIRS hyperscanning experiment with 142 dyads to examine how interpersonal closeness and social interactivity levels affect interpersonal neural synchrony. To manipulate interpersonal closeness, we recruited dyads from three relationship types: close friends, romantic partners, and mother-child pairs. Additionally, each dyad participated in an experimental session involving social interactions at varying levels of interactivity. Specifically, they engaged in three conditions: video co-exposure (passive condition, no direct interaction), rules-based cooperative game (active condition, structured interaction), and free interaction (active condition, unstructured interaction). Based on previous research (De Felice et al., 2024a; Feldman, 2017), we hypothesized that both greater interpersonal closeness (mother-child *>* romantic partners *>* close friends) and higher levels of social interactivity (free interaction *>* cooperative game *>* video co-exposure) would be associated with increased interpersonal neural synchrony between participants.

To test these hypotheses, we initially examined patterns of neural synchrony across different brain regions and compared real dyads to surrogate, non-interacting dyads. Our results revealed significantly higher levels of interpersonal neural synchrony in true dyads compared to surrogate (noninteracting) pairs, across all brain region combinations involving the bilateral IFG and TPJ. This finding aligns with evidence from fNIRS hyperscanning studies in cooperative settings showing statistically significant interpersonal neural synchrony in frontal and temporoparietal regions (Czeszumski et al., 2022). Notably, in our study, the strongest effect when comparing data from real and surrogate dyads was observed in neural synchrony involving the right IFG. The IFG is a key region of the mirror neuron system (Aziz-Zadeh et al., 2006), which plays a crucial role in action understanding and imitation by enabling the brain to simulate observed actions (Aziz-Zadeh et al., 2006; Carr et al., 2003; Iacoboni, 2005; Lyons et al., 2006). Mirror neurons are thought to establish a bridge between passive action observation and social perception encoding (Lyons et al., 2006). A meta-analysis of 485 functional imaging studies on the IFG conducted by Liakakis et al. (2011) identified four functionally distinct clusters in this brain region. In the left hemisphere, one cluster was linked to empathy, another to semantic and phonological processing, and a third to working memory. A fourth cluster, associated with fine movement control, was located in the right hemisphere. Regarding the right IFG, the studies by Adolfi et al. (2017); Bzdok et al. (2012); Hartwigsen et al. (2019) showed that higher-level social cognitive processing occurs in the anterior portions of the right IFG. Similarly, the TPJ is a key hub for mentalizing and perspective-taking, allowing individuals to infer the thoughts, beliefs, and intentions of others (Gallagher et al., 2000; Samson et al., 2004; Van Overwalle, 2009; Yang et al., 2020; Wu et al., 2025). Consistent with the role of these brain regions, Yang et al. (2020) found that neural synchrony within the right dorsolateral prefrontal cortex and within the right TPJ increases among triad members after in-group bonding, with right prefrontal synchrony further predicting stronger impulsivity and hostility toward the out-group. Overall, a possible mechanism supporting the effective processing and facilitation of social exchanges is the temporal alignment of neural activity within mirroring and mentalizing networks to the dynamics of the interaction. This alignment may, in turn, lead to the synchronization of neural signals between individuals, fostering interpersonal neural synchrony and enhancing prediction, communication, and social affiliation (Hoehl et al., 2021).

Building on these findings, we further examined how interpersonal closeness and social interactivity modulate neural synchrony across different brain region combinations. When conducting the analyses on all possible brain region combinations altogether, we observed that both interpersonal closeness and social interactivity influence interpersonal neural synchrony. However, contrary to our hypotheses, close friends and romantic partners exhibited significantly higher synchrony than mother-child dyads. We also observed a non-significant but qualitative trend suggesting that romantic partners showed lower synchrony than close friends. Similarly, in contrast to our expectations, passive social interactions (video co-exposure) elicited higher synchrony than both active, structured interactions (rules-based cooperative game) and active, unstructured interactions (free exchange), particularly among close friends and romantic partners.

The pattern in interpersonal closeness effects persisted when analyzing specific brain region pairs, all involving the IFG. As already observed on physiological synchrony by Bizzego et al. (2019a), interpersonal closeness appears to sometimes have a negative association with synchrony, with more distant compared to closer interpersonal relationships being supported by stronger bio-behavioral synchrony, as long as a meaningful pre-existing relationship is present.. As suggested by Bizzego et al. (2019a), the absence of novelty and the continuous exposure characteristic of highly close relationships may account for this pattern of results. In such established relationships, partners may rely less on physiological or cortical coordination within mirroring and mentalizing networks, as they do not need to rapidly establish a connection or coordinate with each other’s communicative signals under conditions of greater uncertainty. Following Nguyen et al. (2024), high levels of neural synchrony may sometimes reflect compensatory mechanisms to overcome challenges in behavioral coordination. Together with interpersonal closeness, developmental differences may contribute to our findings (Hoehl et al., 2025). In fact, the neural rhythms and functional dynamics of children’s brains might not yet be fully comparable to those of adults (Meng et al., 2022). Brain regions involved in social cognition, such as the IFG and TPJ, continue to mature well into adolescence and early adulthood (Gweon et al., 2012; Kolk and Rakic, 2022; Wang et al., 2020). This prolonged development may affect the efficiency and coordination of neural activity during social interactions, leading to differences in interpersonal neural synchrony between children and adults. Moreover, myelination, synaptic pruning, and changes in connectivity across cortical networks influence the way information is processed and integrated over time. As a result, the mother-child dyads in our study may exhibit lower neural synchrony not necessarily due to interpersonal closeness but rather due to ongoing maturation of brain mechanisms that support shared attention, perspective-taking, and interpersonal attunement in children.

Finally, we observed distinct patterns across different brain region pairs regarding the effect of social interactivity on interpersonal neural synchrony. While some regions (i.e., right IFG–right IFG, right IFG–left TPJ, left TPJ– left TPJ, left TPJ–right TPJ) followed the previously described trend–showing the highest synchrony during video co-exposure, followed by the cooperative game and the lowest during free interaction–other regions (i.e., left IFG– left IFG, left IFG–right TPJ) displayed the opposite pattern, with cooperative game interactions eliciting the highest synchrony, followed by video coexposure, and the lowest synchrony occurring during free interaction. These diverging trends suggest that different neural mechanisms may be engaged depending on the nature of the interaction. In brain regions where synchrony peaks during video co-exposure (primarily the combinations involving the right IFG and the left TPJ), neural alignment may be predominantly driven by shared sensory input in a social context, as both participants passively experience the same audiovisual stimuli. In contrast, regions where the cooperative game leads to the highest synchrony (primarily the combinations involving the left IFG and the right TPJ) may be more involved in joint action coordination, strategic thinking, and shared goal pursuit, processes that require active engagement and real-time behavioral alignment. This pattern is consistent with previous studies on cooperation during Jenga, where the IFG showed statistically significant interpersonal neural synchrony (e.g., Li et al. 2021; Liu et al. 2016), and with intracranial recording studies reporting coordination-related activity in the TPJ during cooperative tasks (Wang et al., 2025). Our results extend these findings by highlighting lateralized effects, with synchrony particularly observed between the left IFG and right TPJ during the cooperative game.

## 5. Limitations

One consideration in our study is the slight variation in data collection methods and sites across participant groups. Specifically, data from adult-adult dyads and mother-child interactions were collected in different countries (Italy and Austria, respectively). While these regions are geographically close and culturally similar, minor contextual or cultural differences cannot be fully excluded and should be considered when interpreting the results. In addition, the two groups were recorded with different fNIRS devices, which were different versions from the same manufacturer. These methodological differences were due to the specific expertise and available equipment of each participating research lab, and are unlikely to compromise the overall validity of our findings. Potentially, such differences can introduce intrinsic variability in raw brain signal recordings (Bizzego et al., 2024), but to date no systematic differences have been reported in the characteristics of signals after standard pre-processing steps have been applied. In our study, to minimize variability related to data collection and laboratory practices, the same researcher (AC) conducted and supervised all sessions at both sites using identical experimental procedures. Additionally, we applied a conservative pre-processing pipeline in order to minimize device-related variance.

Another limitation is the absence of fine-grained behavioral coding. Although our paradigm controls behavior at the macro level through task interactivity, we did not link neural synchrony to moment-to-moment interpersonal behaviors. Such analyses could help clarify how specific actions or interactional dynamics modulate neural synchrony, but they also pose theoretical and methodological challenges (e.g., defining behavioral indices, selecting appropriate synchrony windows). Future studies incorporating detailed behavioral annotations will be an important step to address this point.

Related to group composition, we also acknowledge that age, sex distribution, and relational factors differed substantially between mother–child and adult–adult dyads. In large-scale studies of interpersonal neural synchrony, where dyads rather than individuals are the unit of analysis, it is inherently difficult to achieve perfect matching across groups. Variables such as age, personality traits, or attachment styles are deeply intertwined with the relational context itself and cannot be fully equated across naturally occurring relationships of different types. In designing the present study, we prioritized comparability where feasible.

A further consideration is that our study focused exclusively on the bilateral IFG and TPJ, without monitoring other brain areas, such as the medial prefrontal cortex, which are also relevant for social cognition and interpersonal synchrony. This constraint was primarily due to the number of available fNIRS optodes, which limited our ability to capture a more comprehensive neural representation of social interactions. Future research could benefit from broader brain coverage to provide a more complete understanding of the neural mechanisms underlying interpersonal synchrony.

Lastly, while our findings suggest that the maturation of brain regions involved in social processing may also account for differences across relationship types, including non-parental adult–child dyads and child–child friend dyads in the sample would have allowed for a more precise disentanglement of these effects. Future research could extend the present design to these groups in order to better isolate the contribution of developmental stage and interpersonal closeness to interpersonal neural synchrony.

## 6. Conclusion

In the current study, we employed an fNIRS hyperscanning approach to investigate how interpersonal closeness and social interactivity modulate interpersonal neural synchrony, analyzing data from a diverse sample of 284 participants. To vary interpersonal closeness, we recruited close friends, romantic partners, and mother-child dyads, while social interactivity was manipulated through three conditions: a video co-exposure, a rules-based cooperative game, and a free interaction. Our results highlight that interpersonal neural synchrony exhibited the most pronounced statistical differences when computed between one participant’s right IFG and the other’s frontotemporal regions. Additionally, we identified a robust trend across multiple levels of analysis, indicating that neural synchrony diminishes with interpersonal closeness and potentially with variations in the developmental stage. Furthermore, social interactivity levels had a small effect on interpersonal neural synchrony. Particularly, video co-exposure elicited the strongest synchrony across brain regions, while the cooperative game enhanced synchrony between the left IFG and the other person’s left IFG and right TPJ, underscoring the role of active social interactions in shaping neural alignment in regions involved in the mirror neurons system and supporting theory of mind processes.

These findings contribute to the growing evidence that interpersonal neural synchrony reflects the dynamics of social bonds and shared experiences (Carollo et al., 2025; Lim et al., 2024a,b), with potential implications for developmental and social neuroscience. Understanding how different types of relationships and interactive contexts shape interpersonal neural synchrony may provide valuable insights into social cognition, cooperative behavior, and even therapeutic approaches for individuals with social challenges (e.g., autism spectrum disorder). Future research should explore how individual differences in personality or social skills, as well as relationship quality, further modulate neural synchrony. Furthermore, by comparing dyads and interactive contexts, this study contributes to future hypothesis-driven hyperscanning research, supporting more systematic investigations of the neural mechanisms underlying human social interactions. This study reinforces the idea that human connection is deeply embedded in and supported by behavioral, physiological, and neural mechanisms.

## Authors contribution

Conceptualization: AC, GE; Methodology: AC, AB; Formal analysis: AC, AB; Investigation: AC, VS; Data Curation: AC, AB; Writing – Original Draft: AC; Writing – Review & Editing: AC, AB, VS, CP, SH, GE; Supervision: SH, GE. All authors have read and agreed to the published version of the manuscript.

## Supplementary Materials

**Supplementary Material 1**: Functional Near-Infrared Spectroscopy Cap. **Supplementary Material 2**: Analysis on Deoxygenated Hemoglobin.

## Data and code availability statement

The data and code used for the present study are available upon request to the corresponding author.

## Conflict of interests

The authors declare no conflict of interest.

## Funding

The authors received no specific funding for this work.

## Notes

### Competing Interest Statement

The authors have declared no competing interest.

### Summary of Updates

Added interaction analysis; Added literature on cooperative tasks; Discussed the limitations of the work in more detail;

