## Supplementary Materials 1, 2 for "Interpersonal Neural Synchrony Across Levels of Interpersonal Closeness and Social Interactivity"

### 1. Functional Near-Infrared Spectroscopy Cap

Figure S1 displays the functional near-infrared spectroscopy (fNIRS) cap setup with the optodes arrangement.

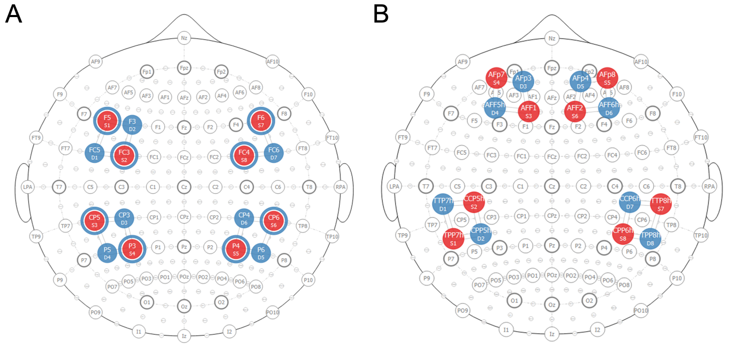


Figure 1: Diagram of the functional near-infrared spectroscopy (fNIRS) cap setup, illustrating the arrangement of optodes for data collection. Solid red circles represent the light sources, while solid blue circles indicate the light detectors. The blue circles surrounding the light sources represent short-separation channels. **A.** Diagram of the cap setup used for close friends and romantic partner dyads in Italy. **B.** Diagram of the cap setup used for mother-child dyads in Austria.

### 2. Analysis on Deoxygenated Hemoglobin

#### *2.1. Neural Synchrony in True Versus Surrogate Dyads*

Table S1 presents the results with the comparison of interpersonal neural synchrony between true, interacting and surrogate, non-interacting dyads. All combinations of regions of interest show a statistically significant effect, with true dyads showing higher levels of interpersonal neural synchrony as compared to surrogate dyads ($q < .001$).

#### *2.2. Interpersonal Closeness and Social Interactivity on Neural Synchrony (All Combinations of Regions)*

We used a linear mixed model to assess the impact of interpersonal closeness and social interactivity on interpersonal neural synchrony across all regions of interest. Our findings reveal that social interactivity had a statistically significant effect on synchrony ($F(2,\ 71{,}598) = 1755.52$, $p < .001$), whereas interpersonal closeness did not ($F(2,\ 131) = 1.52$, $p > .05$). Additionally, we observed an interaction effect of interpersonal closeness and

| Region A | Region B | Estimate | Standard Error | $t$-value | $q$-value |
|---|---|---|---|---|---|
| Left IFG | Left IFG | 0.004 | 0.0002 | 15.17 | $< .001$ *** |
| Left IFG | Right IFG | 0.004 | 0.0002 | 20.60 | $< .001$ *** |
| Left IFG | Left TPJ | 0.003 | 0.0002 | 20.39 | $< .001$ *** |
| Left IFG | Right TPJ | 0.003 | 0.0002 | 13.18 | $< .001$ *** |
| Right IFG | Right IFG | 0.005 | 0.0003 | 18.70 | $< .001$ *** |
| Right IFG | Left TPJ | 0.004 | 0.0002 | 22.07 | $< .001$ *** |
| Right IFG | Right TPJ | 0.003 | 0.0002 | 12.76 | $< .001$ *** |
| Left TPJ | Left TPJ | 0.005 | 0.0003 | 18.20 | $< .001$ *** |
| Left TPJ | Right TPJ | 0.003 | 0.0002 | 15.63 | $< .001$ *** |
| Right TPJ | Right TPJ | 0.004 | 0.0004 | 8.77 | $< .001$ *** |

Table 1: Comparison of interpersonal neural synchrony between true and surrogate dyads. For each combination of region of interest (i.e., inferior frontal gyrus (IFG) and temporoparietal junction (TPJ)), we report the estimated contrast between true and surrogate dyads, along with the standard error, $t$-value, and corrected $q$-value. (*** $q < .001$).

social interactivity on interpersonal neural synchrony ($F(4,$ 71,600$) = 75.67$, $p < .001$).

*Post-hoc* analyses showed significant differences in interpersonal neural synchrony between video co-exposure and rules-based cooperative game (estimate = -0.008, SE = 0.000126, $z$ = -64.115, $q < .001$), video co-exposure and free verbal interaction (estimate = -0.004, SE = 0.000124, $z$ = -30.916, $q < .001$), and rules-based cooperative game and free verbal interaction (estimate = 0.004, SE = 0.000126, $z$ = 33.86, $q < .001$). This pattern was consistent across levels of interpersonal closeness (see Table 2). Unlike the analyses on HBO, the results for HBR reveal that only the level of social interactivity has a significant effect on interpersonal neural synchrony, with no influence of interpersonal closeness. Specifically, the cooperative rules-based game condition exhibited the highest synchrony levels, suggesting that structured social interactions may enhance neural alignment more effectively than passive co-experiences or unstructured verbal exchanges.

#### *2.3. Interpersonal Closeness and Social Interactivity on Neural Synchrony (Individual Combinations of Regions)*

All linear mixed models conducted on data from individual combinations of regions of interest revealed a statistically significant effect of social interactivity, but not social closeness, on neural synchrony. Particularly, this pattern of results was observed in the following combinations of regions of interest:

| **Contrast** | **Estimate** | **Standard Error** | ***z*-ratio** | ***q*** |
|---|---|---|---|---|
| | | **Close friends** | | |
| Video–Cooperative game | -0.00878 | 0.0002 | -48.67 | < .001 *** |
| Video–Free interaction | -0.00376 | 0.0002 | -21.47 | < .001 *** |
| Cooperative game–Free interaction | 0.00502 | 0.0002 | 27.92 | < .001 *** |
| | | **Romantic partners** | | |
| Video–Cooperative game | -0.00976 | 0.0002 | -40.17 < .001 *** | |
| Video–Free interaction | -0.00376 | 0.0002 | -15.54 | < .001 *** |
| Cooperative game–Free interaction | 0.00600 | 0.0002 | 24.55 | < .001 *** |
| | | **Mother-child** | | |
| Video–Cooperative game | -0.00497 | 0.0003 | -19.36 | < .001 *** |
| Video–Free interaction | -0.00402 | 0.0003 | -15.70 | < .001 *** |
| Cooperative game–Free interaction | 0.00095 | 0.0003 | 3.78 | .006 ** |
| | | **Video co-exposure** | | |
| Close friends–Romantic partners | 0.00010 | 0.0011 | 0.09 | > .05 |
| Close friends–Mother-child | -0.00323 | 0.0012 | -2.65 | > .05 |
| Romantic partners–Mother-child | -0.00334 | 0.0014 | -2.44 | > .05 |
| | | **Cooperative game** | | |
| Close friends–Romantic partners | -0.00088 | 0.0011 | -0.77 | > .05 |
| Close friends–Mother-child | 0.00058 | 0.0012 | 0.47 | > .05 |
| Romantic partners–Mother-child | 0.00146 | 0.0014 | 1.07 | > .05 |
| | | **Free interaction** | | |
| Close friends–Romantic partners | 0.00011 | 0.0011 | 0.09 | > .05 |
| Close friends–Mother-child | -0.00349 | 0.0012 | -2.86 | > .05 |
| Romantic partners–Mother-child | -0.00360 | 0.0014 | -2.64 | > .05 |

Table 2: Results from the *post-hoc* pairwise comparisons of the interaction between interpersonal closeness and social interactivity on interpersonal neural synchrony (across all brain region combinations). For each level of one factor, the table reports the estimated contrasts between two levels of the other factor, together with the standard error, $z$-ratio, and corrected $q$-value. (** $q < .01$; *** $q < .001$).

left IFG–left IFG (interpersonal closeness: $F(2,\ 131.90) = 0.29$, $p > .05$, $q > .05$; social interactivity: $F(2,\ 5581.50) = 176.62$, $p < .001$, $q < .001$), left IFG–right IFG (interpersonal closeness: $F(2,\ 131.70) = 1.53$, $p > .05$, $q > .05$; social interactivity: $F(2,\ 11{,}070.90) = 360.45$, $p < .001$, $q < .001$), left IFG–left TPJ (interpersonal closeness: $F(2,\ 131.80) = 1.74$, $p > .05$, $q > .05$; social interactivity: $F(2,\ 11{,}003.40) = 399.04$, $p < .001$, $q < .001$), left IFG–right TPJ (interpersonal closeness: $F(2,\ 128.50) = 0.45$, $p > .05$, $q > .05$; social interactivity: $F(2,\ 6597.50) = 236.02$, $p < .001$, $q < .001$), right IFG–right IFG (interpersonal closeness: $F(2,\ 130.50) = 0.73$, $p > .05$, $q > .05$; social interactivity: $F(2,\ 5387.50) = 99.99$, $p < .001$, $q < .001$), right IFG–left TPJ (interpersonal closeness: $F(2,\ 132.10) = 1.52$, $p > .05$, $q > .05$; social interactivity: $F(2,\ 10{,}828.30) = 328.79$, $p < .001$, $q < .001$), right IFG–right TPJ (interpersonal closeness: $F(2,\ 126.20) = 1.28$, $p > .05$, $q > .05$; social interactivity: $F(2,\ 6448.60) = 183.36$, $p < .001$, $q < .001$), left TPJ–left TPJ (interpersonal closeness: $F(2,\ 129.50) = 1.19$, $p > .05$,

$q > .05$; social interactivity: $F(2, 5330.50) = 138.60$, $p < .001$, $q < .001$), left TPJ–right TPJ (interpersonal closeness: $F(2, 127.90) = 2.61$, $p > .05$, $q > .05$; social interactivity: $F(2, 6423.70) = 233.50$, $p < .001$, $q < .001$), and right TPJ–right TPJ (interpersonal closeness: $F(2, 121.32) = 0.95$, $p > .05$, $q > .05$; social interactivity: $F(2, 2013.09) = 37.54$, $p < .001$, $q < .001$).

Additionally, pairwise comparisons of marginal means indicated that, across all individual combinations of regions of interest, interpersonal neural synchrony was highest during the rules-based cooperative game, followed by free interaction, and lowest during video co-exposure (see Table S3 for a summary of the results).

| **Contrast** | **Estimate** | **Standard Error** | **z-ratio** | ***q*** |
|---|---|---|---|---|
| **Left inferior frontal gyrus – left inferior frontal gyrus** | | | | |
| Video–Cooperative game | -0.00784 | 0.0004 | -18.43 | < .001 *** |
| Video–Free interaction | -0.00254 | 0.0004 | -6.06 | < .001 *** |
| Cooperative game–Free interaction | 0.00530 | 0.0004 | 12.57 | < .001 *** |
| **Left inferior frontal gyrus – right inferior frontal gyrus** | | | | |
| Video–Cooperative game | -0.00830 | 0.0003 | -26.85 | < .001 *** |
| Video–Free interaction | -0.00408 | 0.0003 | -13.43 | < .001 *** |
| Cooperative game–Free interaction | 0.00421 | 0.0003 | 13.73 | < .001 *** |
| **Left inferior frontal gyrus – left temporoparietal junction** | | | | |
| Video–Cooperative game | -0.00875 | 0.0003 | -28.25 | < .001 *** |
| Video–Free interaction | -0.00436 | 0.0003 | -14.29 | < .001 *** |
| Cooperative game–Free interaction | 0.00439 | 0.0003 | 14.23 | < .001 *** |
| **Left inferior frontal gyrus – right temporoparietal junction** | | | | |
| Video–Cooperative game | -0.00837 | 0.0004 | -21.71 | < .001 *** |
| Video–Free interaction | -0.00438 | 0.0004 | -11.54 | < .001 *** |
| Cooperative game–Free interaction | 0.00398 | 0.0004 | 10.41 | < .001 *** |
| **Right inferior frontal gyrus – right inferior frontal gyrus** | | | | |
| Video–Cooperative game | -0.00665 | 0.0005 | -14.11 | < .001 *** |
| Video–Free interaction | -0.00291 | 0.0005 | -6.28 | < .001 *** |
| Cooperative game–Free interaction | 0.00374 | 0.0005 | 7.98 | < .001 *** |
| **Right inferior frontal gyrus – left temporoparietal junction** | | | | |
| Video–Cooperative game | -0.00835 | 0.0003 | -25.60 | < .001 *** |
| Video–Free interaction | -0.00365 | 0.0003 | -11.335 | < .001 *** |
| Cooperative game–Free interaction | 0.00471 | 0.0003 | 14.48 | < .001 *** |
| **Right inferior frontal gyrus – right temporoparietal junction** | | | | |
| Video–Cooperative game | -0.00799 | 0.0004 | -19.13 | < .001 *** |
| Video–Free interaction | -0.00425 | 0.0004 | -10.32 | < .001 *** |
| Cooperative game–Free interaction | 0.00375 | 0.0004 | 9.02 | < .001 *** |
| **Left temporoparietal junction – left temporoparietal junction** | | | | |
| Video–Cooperative game | -0.00767 | 0.0005 | -16.56 | < .001 *** |
| Video–Free interaction | -0.00307 | 0.0005 | -6.71 | < .001 *** |
| Cooperative game–Free interaction | 0.00461 | 0.0005 | 9.96 | < .001 *** |
| **Left temporoparietal junction – right temporoparietal junction** | | | | |
| Video–Cooperative game | -0.00859 | 0.0004 | -21.51 | < .001 *** |
| Video–Free interaction | -0.00491 | 0.0004 | -12.49 | < .001 *** |
| Cooperative game–Free interaction | 0.00367 | 0.0004 | 9.22 | < .001 *** |
| **Right temporoparietal junction – right temporoparietal junction** | | | | |
| Video–Cooperative game | -0.00619 | 0.0007 | -8.66 | < .001 *** |
| Video–Free interaction | -0.00312 | 0.0007 | -4.42 | < .001 *** |
| Cooperative game–Free interaction | 0.00307 | 0.0007 | 4.31 | .001 ** |

Table 3: Results of the *post-hoc* pairwise comparisons examining the effect of social interactivity levels on interpersonal neural synchrony across different brain region combinations. For each combination, we report the estimated contrast between two levels of social interactivity, along with the standard error, $z$-ratio, and corrected $q$-value. (** $q < .01$; *** $q < .001$).
